# Structural basis of bifunctional CTP/dCTP synthase

**DOI:** 10.1101/2023.02.19.529158

**Authors:** Chen-Jun Guo, Zherong Zhang, Jiale Zhong, Ji-Long Liu

## Abstract

Nucleotides are important molecules of life. According to the sugar, nucleotides can be divided into nucleotides and deoxynucleotides, which are the basic components of RNA and DNA respectively. CTP synthase (CTPS) catalyzes the last step in the de novo synthesis of CTP, which can form cytoophidia in cells from all three domains of life. We have recently found that CTPS combines with NTPs to form filaments, and solved the structures of CTPS bound with NTPs. Previous biochemical studies have shown that CTPS can also serve as dCTPS, i.e. CTPS/dCTPS can not only bind UTP, ATP and GTP to generate CTP, but also bind deoxynucleotides to generate dCTP. However, the structural basis of the bifunctional enzyme CTPS/dCTPS binding deoxynucleotide is not clear. In this study, we find that CTPS/dCTPS can form filaments bound with deoxynucleotides. Biochemically, we compare the binding and reaction characteristics of the corresponding nucleotides/deoxynucleotides and CTPS/dCTPS. Using cryo-electron microscopy, we solve the the structure of CTPS/dCTPS bound with deoxynucleotides at near-atomic resolution. This study not only provides a structural basis for understanding the catalysis and regulation of bifunctional CTPS/dCTPS, but also opens a door for further exploration the compartmentation of CTPS/dCTPS inside a cell.

## Main

Cytidine triphosphate synthase (CTPS), responsible for the final step of de novo synthesis of cytidine triphosphate(CTP), plays a critical role in the metabolic processes of living system due to the irreplaceable effect of its product CTP in DNA, RNA, and phospholipid synthesis[1–4]. CTPS is upregulated in cancer, and is thought to be an attractive target for cancer[5–12] and immunosuppression[13, 14] as well as for protozoal[15], viral[16] and Mycobacterium tuberculosis infections[17–19].

CTPS compromises two domains, the N-terminal ammonia ligase (AL) domain and the C-terminal glutamine amide transferase (GAT) domain. UTP is phosphorylated by hydrolyzing ATP to active state in AL domain. The intermediate product 4Pi-UTP then be ammoniated to product CTP by ammonia generated by glutamine hydrolyzing in GAT domain[3, 20–26]. In the whole reaction, allosteric regulator GTP synchronizes two independent domains and plays a key role in ammonia tunnel regulation[20, 22–24, 26–33]. The regulation of CTPS by its ligands has been explored for decades through different experimental methods. In the beginning, the unique regulation of ligands such as GTP was explored through a variety of beautiful biochemical experiments[1, 2, 5, 8, 20–24, 27, 34, 35], and finally the binding sites of ligands are determined by means of X-ray crystallography and cryogenic electron microscopy[14, 33, 36–40]. In addition, CTPS is also regulated by post-translational modifications such as phosphorylation, and has been verified for different species[41–45].

Previous study reveals that CTPS can form filamentous structure termed ‘cytoophidia’ under specific condition which varies in different species in prokaryotic and eukaryotic cells[46–56]. The filamentation of CTPS protein with substrates or products in vitro also provides a basis for the compartmentation in vivo[13, 14, 38–40, 55, 57]. By combining structural study and kinetic assay, people found the regulation of CTPS by filamentation is dedicate and diverse.

Deoxynucleotide triphosphates(dNTPs) are the precursor for the DNA synthesis, play key roles in DNA replication, recombination, and repair in cells. Ribonucleotide reductase (RNR) are thought to catalyze the four ribonucleotide diphosphates(rNDPs) to the respective deoxynucleotide diphosphates (dNDPs), and then convert to dNTP[58]. In 1999, carman found a potential way that CTPS can also catalyzes such biosynthesis of dCTP[59]. However, the structural basis for this interesting phenomenon remains unclear.

Using Cryo-EM, here we successfully determine the structure of CTPS/dCTPS with deoxyribonucleotides and inhibitor 6-diazo-L-noreleucine (DON). By using 3D variability analysis, multiple states of CTPS/dCTPS are separated, providing a systematic structural basis for CTPS/dCTPS catalysis. Combined with kinetic assay and other means, we find that 1) dATP is comparable to ATP in its role as energy supplier; 2) dUTP is a slower substrate for the reaction; 3) dGTP is a much gentler allosteric regulator than GTP; and 4) dCTP is a strong inhibitor for CTPS/dCTPS. Based on these results, we propose a bifunctional diagram for CTPS/dCTPS.

## Results

### Determination of CTPS/dCTPS dynamic structure under deoxy-substrate condition

Using Negative-Stain EM, we observed that CTPS/dCTPS mixed with dATP, dUTP and dGTP (referred to as deoxy-substrate conditions below) was capable of forming filaments; the same phenomenon was also observed directly in the Cryo-EM sample (Fig. 1A; Supplementary Fig. S1). This led us to investigate into the role of deoxyribonucleotides in CTPS/dCTPS’ catalysis and their respective structural basis. So, we used Cryo-EM to determine and acquire the structure of CTPS/dCTPS under deoxy-substrate (CTPS/dCTPS^+deo^) conditions at 3.5-Å resolution, referred as Average model from here on (Supplementary Fig. S2). In Average model, CTPS/dCTPS^+deo^ protomers form tetramers (Fig. 1B), which then assembles into filament (Fig. 1C). The filament’s rise is 104 Å, and its twist is 50°. Similar to CTPS/dCTPS in Mix state, CTPS/dCTPS^+deo^ tetramers form filaments through 355H on the tetramer-tetramer interface (Fig. 1D).

**Fig. 1.**
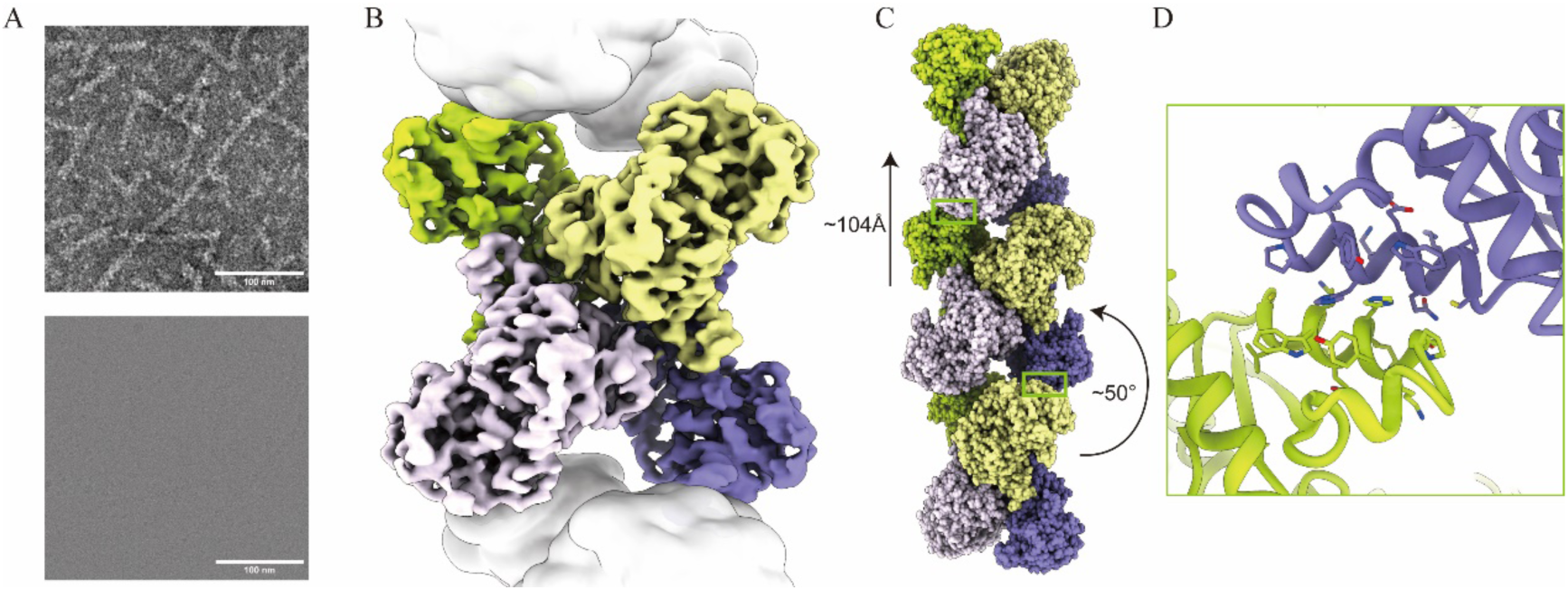
Overall structure of CTPS/dCTPS. A, Negative-stain EM images of 1.764 µM CTPS/dCTPS mixed with 2 mM dATP, 2 mM dUTP, 2 mM dGTP, 0.6 mM DON, 8 mM MgCl_2_ and image of Cryo-EM sample in deoxy-substrate condition. The scale bars are 100 nm as marked in the images. B,3.5Å resolution structure of CTPS/dCTPS tetramer, with each protomer colored differently. Adjacent CTPS/dCTPS tetramers that form filaments with the tetramer in center are indicated by white surface. C, Overall structure of CTPS/dCTPS filament and its helical parameters: rise is 104 Armstrong, and twist is 50 degrees, as indicated. D, Model of the tetramer-tetramer interface of CTPS/dCTPS.

Through the 3D variable analysis, we then categorized our data of dynamic CTPS/dCTPS^+deo^ structure into three states: the “Open” state (Fig. 2A,B), the “Semi-open” state (Fig. 2C,D) and the “Closed” state (Fig. 2E,F). All three states are classified by different stages of the CTPS/dCTPS^+deo^ “wing” structure’s dynamic movement to the binding of dGTP. As their names suggest, “Open” state suggests the beginning of the dGTP binding process; “Semi-open” state corresponds to the preliminary binding of dGTP; and “Closed” state represents the tight binding of dGTP. The conformations also attest to three distinctive states. While “Semi-open” state model is overlaid on the Average state model (Fig. 2D), “Open” state model and “Closed” state model swings to the upper left (Fig. 2B) of and the right (Fig. 2F) of the Average state model, respectively.

**Fig. 2.**
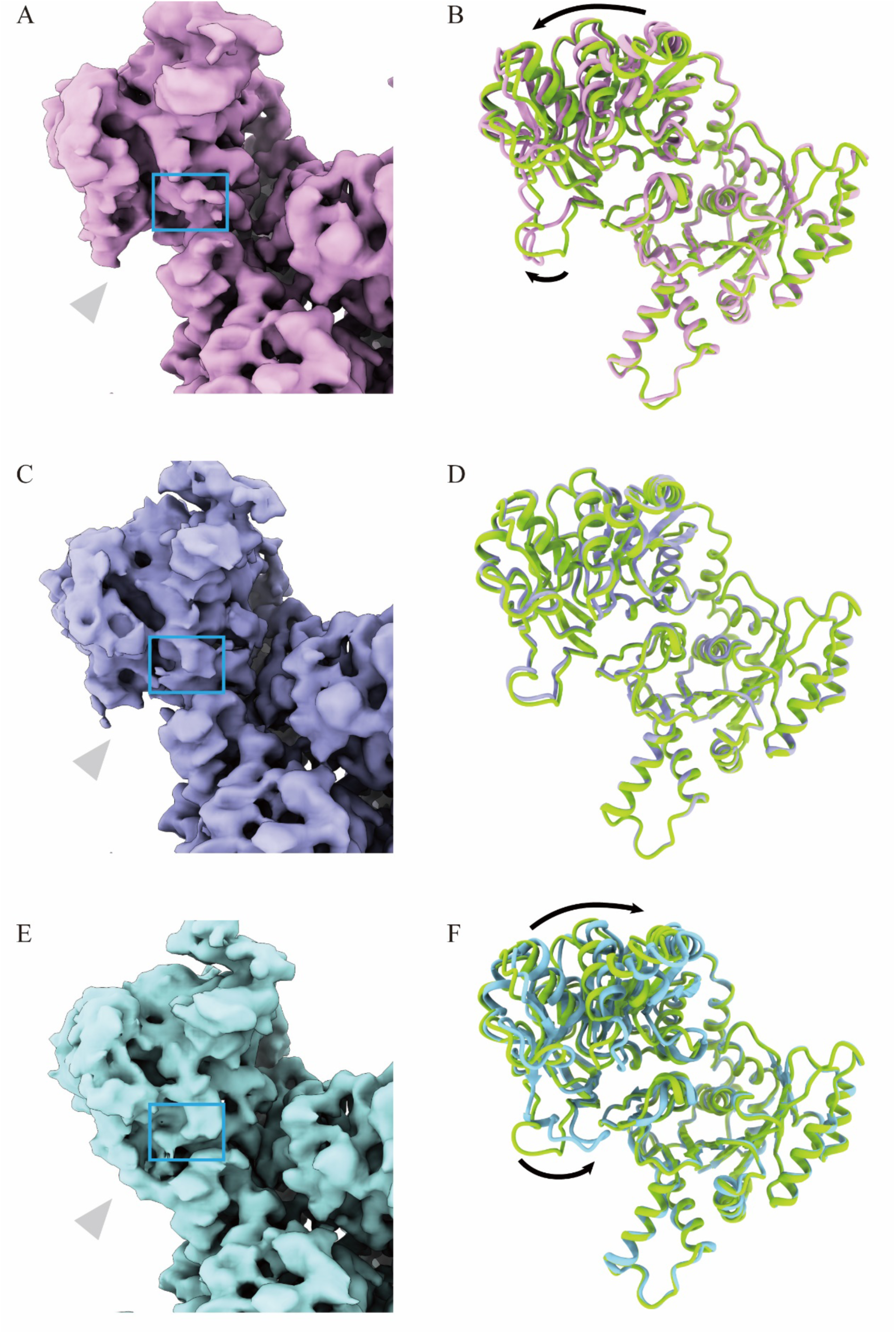
Three states of CTPS/dCTPS in deoxy-substrate condition. A, “Open” state map of CTPS/dCTPS. dGTP density is marked by blue box and the wing density is pointed by grey arrow. B, Structural comparison of CTPS/dCTPS in “open” state (red) and average state (green). C, “Semi-open” state map of CTPS/dCTPS. D, Structural comparison of the CTPS/dCTPS in “Semi-open” state (purple) and average state (green). The models show no significant conformational differences between the two states. E, “Closed” state map of CTPS/dCTPS. dGTP density is marked by blue box. F, Structural comparison of the CTPS/dCTPS in “closed” state (blue) and average state (green). Overall conformational shift to AL domain is indicated by arrows.

The acquisition of these high-resolution models provides the structural basis for that deoxyribonucleotides stimulate filament formation of CTPS/dCTPS. The three distinctive states of CTPS/dCTPS also suggest that deoxyribonucleotides induce conformational switches in CTPS/dCTPS’ catalytic process.

### dATP is comparable to ATP in its role as energy supplier to CTPS/dCTPS catalysis

Adenine and deoxyribose ring are resolved with good density; the triphosphates also demonstrated stable binding with CTPS/dCTPS (Fig. 3A). The deoxyribose ring corresponds with the right amount of density, which validated that the bound ligand is dATP.

**Fig. 3.**
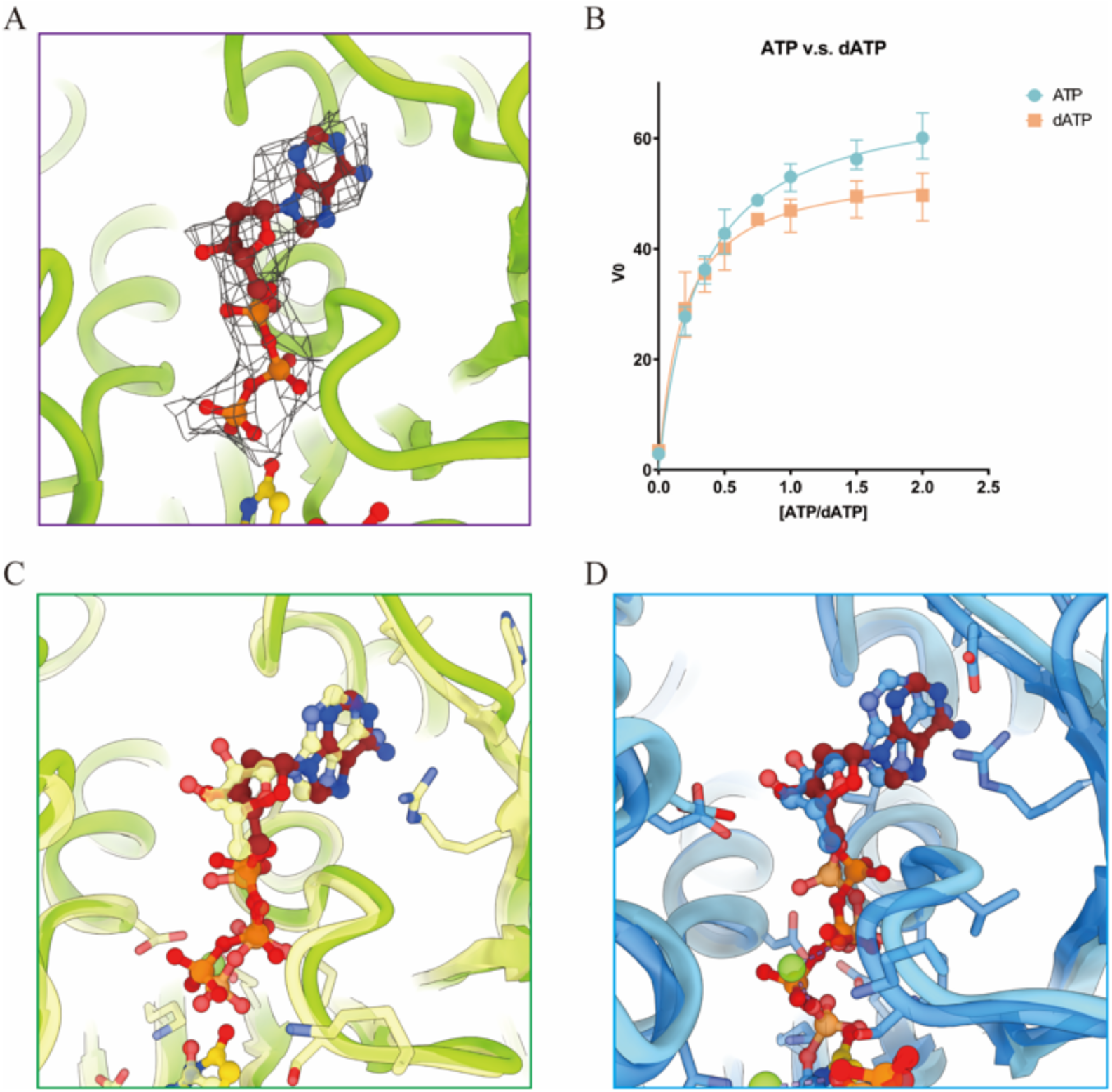
Structural and kinetic characterization of CTPS/dCTPS in both ATP and dATP binding modes. A, dATP binding site in CTPS/dCTPS under deoxy-substrate condition. B, Kinetic curves of CTPS/dCTPS in both ATP and dATP binding modes. X-axis indicates ATP or dATP concentrations. Y-axis indicates the initial velocity of the enzymatic reaction. C, Structural comparison between the ATP binding pocket in Mix state (yellow) and the dATP binding pocket in Average state (green). D, Structural comparison between the ATP binding pocket in DON state (dark blue) and the dATP binding pocket Average state (light blue).

Using a classic Michaelis-Menten equation, we fitted both ATP-driven and dATP-driven kinetic curves (Fig. 3B). ATP curve shows a generally higher V_max_ than dATP curves, which means that CTPS/dCTPS can react at a slightly faster rate using ATP than dATP. Interestingly, CTPS/dCTPS’ binding affinity towards a substrate, as measured by k_cat_/K_m_, indicates that it slightly prefers binding with dATP than ATP, although the preference is relatively marginal. We used classic Michaelis-Menten equation to fit both activities. ATP-driven activity’s V_max_ and K_m_ are 68.59 and 0.3021, respectively. dATP-driven activity’s V_max_ and K_m_ are 55.08 and 0.1818, respectively.

To find structural basis for the kinetic observations, we then compared structures of the ATP binding pocket in Mix state and the dATP binding pocket in Average state (Fig. 3C). Positions of the deoxyribose ring and adenine in Average state did not differ too much from the ribose ring and adenine in Mix state; positions of the alpha and beta phosphates also remain quite similar, which potentially allows breakage of the phosphoanhyride bond. We also compared binding pockets of ATP and dATP in DON state and Closed state, respectively (Fig. 3D). We observed obvious shift in positions of the deoxyribose ring and adenosine from their counterparts in DON state. We also observed that the dATP binding pocket conformation also changed. These structural differences may underlie the difference between the kinetic parameters of ATP-driven enzyme activity and those of dATP-driven enzyme activity.

### dUTP acts a slower substrate for CTPS/dCTPS activity than UTP

We have also located dUTP at its binding pocket, and the ligand is solved with good density (Fig. 4A). We then performed kinetic measurements of CTPS/dCTPS activity using UTP as substrate (UTP activity) and CTPS/dCTPS activity using dUTP as substrate (dUTP activity) (Fig. 4B). The curves showcase a significant difference between CTPS/dCTPS’ binding affinities for each substrate. Due to a lower K_m_ and higher k_cat_/K_m_ of UTP activity than dUTP activity, CTPS/dCTPS prefers binding to UTP than dUTP. Interestingly, we also observed substrate inhibition in both UTP and dUTP activities (Supplementary Fig. S3), which was not explicitly mentioned in previous research.

**Fig. 4.**
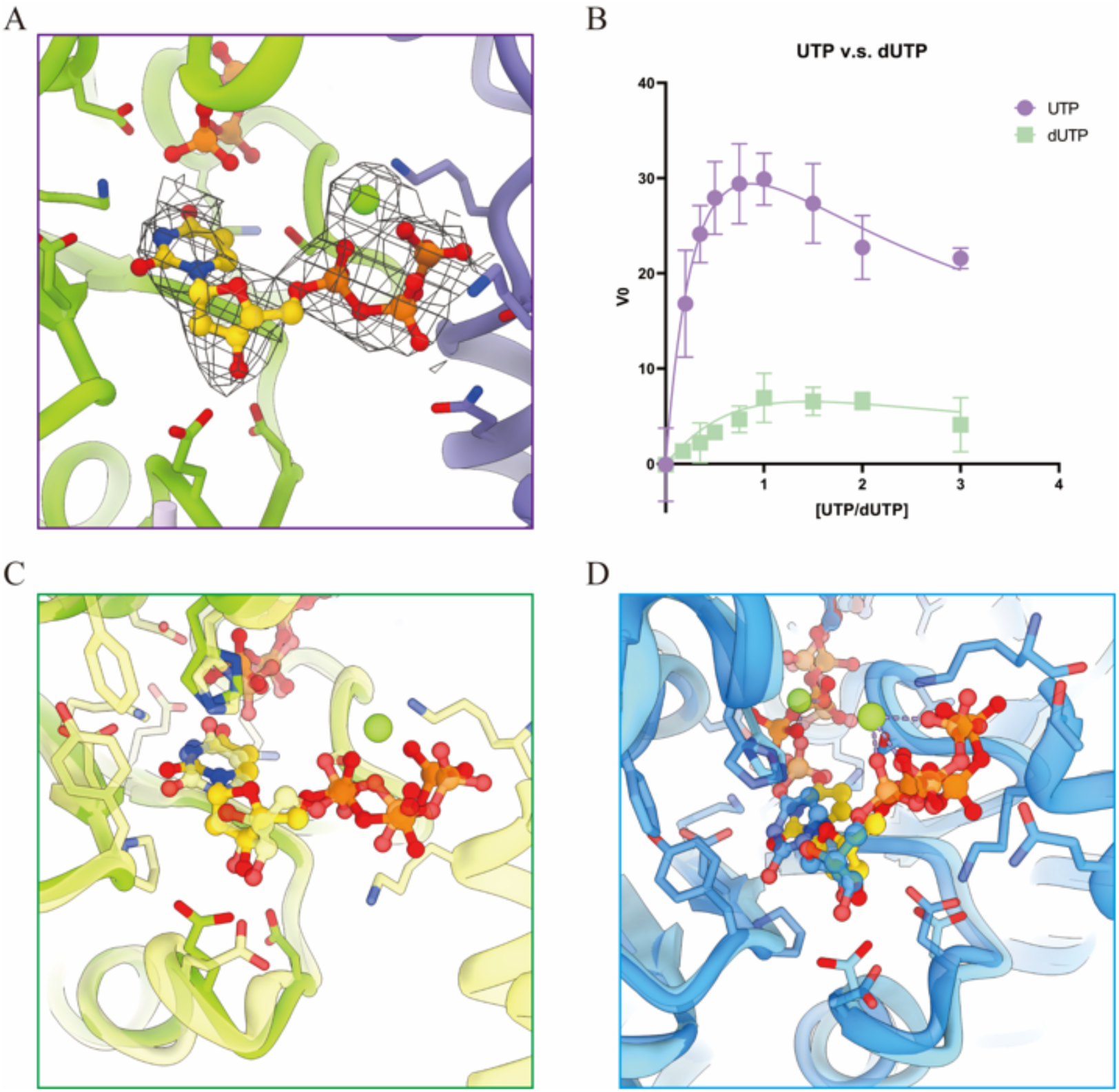
Structural and kinetic characterization of CTPS/dCTPS in both UTP and dUTP binding modes. A, dUTP binding site in CTPS/dCTPS under deoxy-substrate condition. B, Kinetic curves of CTPS/dCTPS activity using UTP and dUTP as substrates. C, Structural comparison between the UTP binding pocket in Mix state (yellow) and the dUTP binding pocket in Average state (green). D, Structural comparison between the UTP binding pocket in DON state (dark blue) and the dUTP binding pocket Average state (light blue).

Using a substrate inhibition model, we deduced that CTPS/dCTPS is more prone to be inhibited by dUTP than UTP because of dUTP activity’s higher K_i_. UTP-driven activity’s V_max_, K_m_ and K_i_ are 60, 0.4512 and 1.668, respectively. dUTP-driven activity’s V_max_, K_m_ and K_i_ are 24.52, 2 and 1.054, respectively. The substrate inhibition equation is as follow:

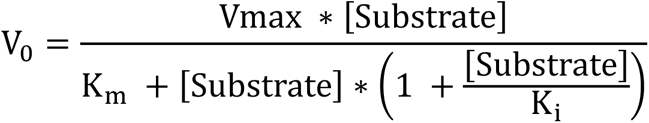

We then proceed to find these catalytic differences’ structural basis. We compared structures of the UTP binding pocket in Mix state and the dUTP binding pocket in Average state (Fig. 4C); we also compared the UTP binding pocket in DON state and the dUTP binding pocket in Average state (Fig. 4D). Both comparisons illustrate that in either state, positions of dUTP’s three chemical groups all shift visibly from positions of their ribonucleotide counterparts. The difference in binding modes may be attributed to dUTP’s interface with amino acids 152D and 154E, which are both highly conserved (Supplementary Fig. S4). It appears that the missing 3’ hydroxyl group on dUTP’s ribose ring results in a weaker interaction between the ribose ring and the two amino acids with negatively charged side chains. These structural and conformational differences provide basis for why CTPS/dCTPS catalyzes dUTP as a substrate but exhibits different kinetic behavior.

### dGTP induces gentler allosteric regulation of CTPS/dCTPS activity than GTP

In the dGTP binding model we resolved, fragmented density impaired a subatomic presentation of the dGTP binding pocket. We overlaid DON state model onto the density of the Average state model, and results showed that ligand density in dGTP binding pocket is similar to that in GTP binding pocket in DON state. Despite that resolution is not optimal, our analysis indirectly provided validation for the presence dGTP in our model (Fig. 5A).

**Fig. 5.**
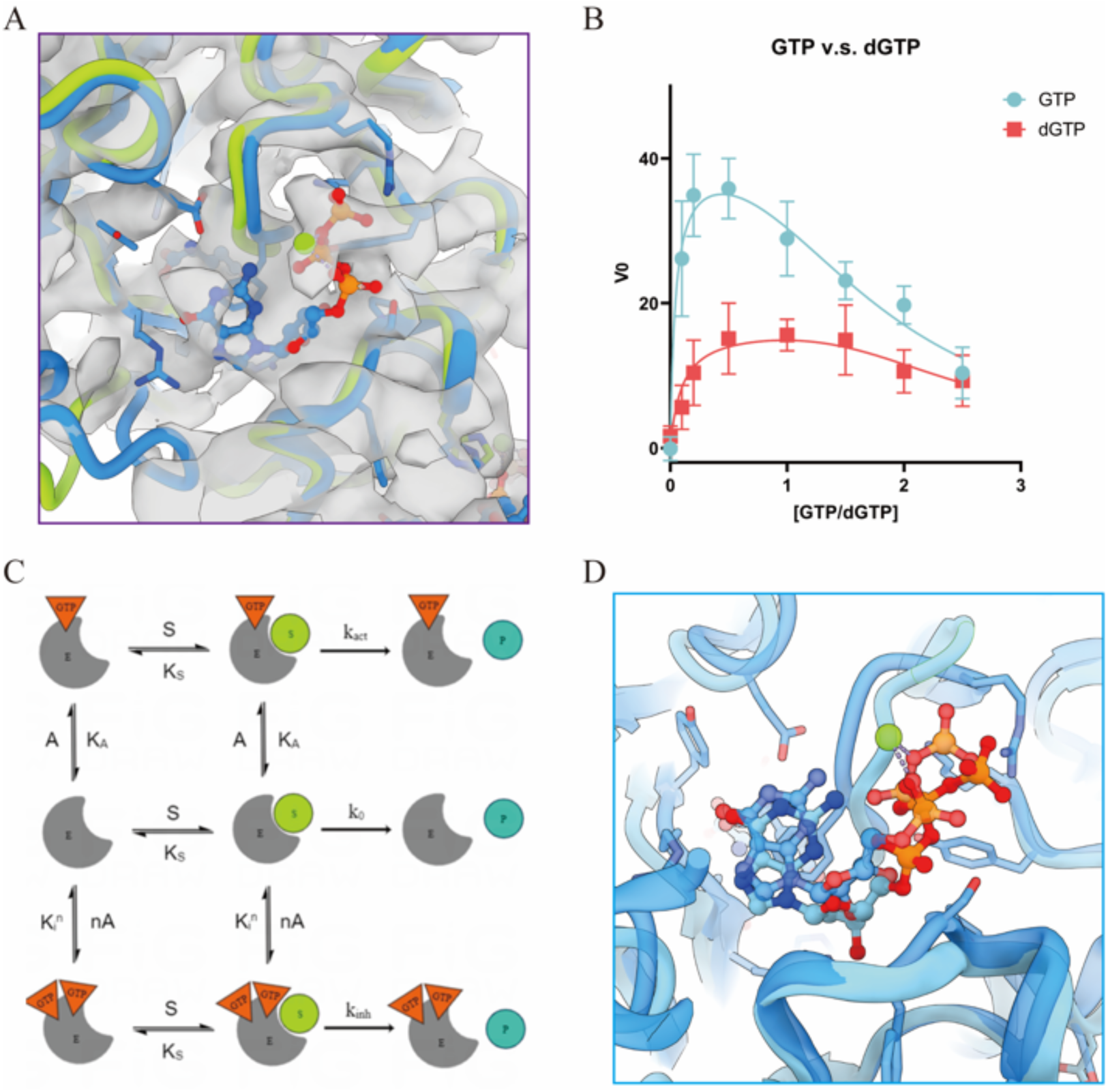
Structural, kinetic, and schematic characterization of CTPS/dCTPS in both GTP and dGTP binding modes. A, Model of dGTP binding pocket in Average state (green) is aligned with model of the GTP binding pocket in DON state (blue), with the average map displayed in transparent grey. B, Kinetic curves of CTPS/dCTPS with the allosteric regulator being either GTP or dGTP. C, Schematic representation of GTP’s different modes of regulation of glutamine-dependent CTP formation in CTPS/dCTPS, modified based on scheme by Bearne [2004 paper]. This scheme is similarly applied to dGTP’s regulation of dCTP formation in CTPS/dCTPS. The central idea of the scheme is that CTPS is capable of forming complex with either one GTP, creating a E·GTP complex, or more than one GTP, which yields E·GTP^n^ complex. E is CTPS/dCTPS; P is CTP or dCTP; A is GTP or dGTP; S is Glutamine; K_A_ is dissociation constant for E·GTP complex; K_S_ is dissociation constant for glutamine; K_i_ is the dissociation constant for E·GTP complex; n is the number of GTP that associates with the enzyme in E·GTP^n^ complex; k_act_, k_0_ and k_inh_ are rates at which CTP or dCTP forms. The two orange triangles on the bottom of the scheme are purely illustrative, so they do not reflect the true numbers (n) of GTP that can form E·GTP^n^ complex. D, Structural comparison between the GTP binding pocket in DON state (dark blue) and the dGTP binding pocket average state (light blue).

We then performed kinetic measurements of CTPS/dCTPS activity when regulated by GTP and dGTP, respectively (Fig. 5B). Considering the intricate behavior GTP exhibited as an allosteric regulator in CTPS activity, we analyzed the kinetic data using the biophysical model created by Bearne et al. (Fig. 5C). This model is similarly applied to dGTP’s regulation of CTPS/dCTPS activity. GTP-driven activity is fitted best to the model when k_0_, k_act_, K_A_, k_inh_, n, K_i_ are 0, 40, 0.04353, 0, 3.276 and 0.5703, respectively. dGTP-driven activity is fitted best to the model when k_0_, k_act_, K_A_, k_inh_, n, K_i_ are 0, 16.78, 0.1, 4.102, 5 and 1.208, respectively. The kinetic model equation is as follows:

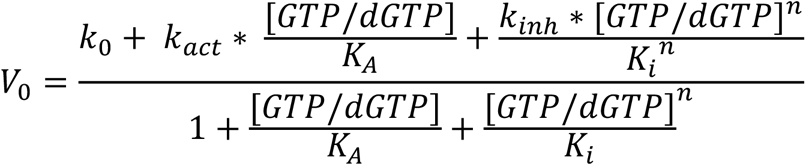

The central idea of the model is that CTPS is capable of forming complex with either one GTP, creating a E·GTP complex, or more than one GTP, which yields E·GTP^n^ complex.

Kinetic data and analysis generated several fascinating insights into catalytic behavior of GTP and dGTP. For most concentrations we have measured, GTP coordinates CTPS/dCTPS activity better than dGTP does, as GTP-driven activity has a higher k_act_ than dGTP-driven activity does. Inferring from dGTP-driven activity’s higher K_A_, dGTP is more likely to dissociate with the enzyme-dGTP complex, which reduces dCTP production in enzyme-dGTP complex relative to CTP production in enzyme-GTP complex. Interestingly, we also found that GTP-driven activity has lower K_i_, which means that GTP is less likely to dissociate from enzyme-multi-GTP complex than dGTP does. According to our fitting, enzyme-multi-GTP complex is unable to catalyze any CTP, which makes GTP’s tendency to stay in such complex an inhibition to CTP production. This aspect of GTP-driven activity partially explains the steep drop in the GTP kinetic curve. On the other hand, dGTP’s higher K_i_ and k_inh_ allow it to maintain a relatively stable dCTP production even when enzyme-multi-dGTP complexes form at higher dGTP concentrations.

To summarize, although dGTP generally does not regulate CTPS/dCTPS activity as well as GTP does, dGTP is a much gentler regulator: it shows a much stable kinetic curve and unlike GTP-driven activity, dGTP-driven activity resists strong inhibition at higher dGTP concentrations.

### Both CTP and dCTP induce filament formation and inhibit CTPS/dCTPS catalytic activity

Carman et al. previously showed that CTPS/dCTPS can tetramerize under dCTP and CTP conditions. To further explore the phenomenon, we used Negative-Stain EM to observe CTPS/dCTPS mixed with dCTP (Fig. 6A). CTPS/dCTPS filaments formed, which suggest that dCTP is capable of inducing filamentation of CTPS/dCTPS. We also performed a set of absorbance readings of CTPS/dCTPS activities to qualitatively verify the inhibitory effects of CTP and dCTP (Fig. 6B). It is quite clear that readings of CTPS/dCTPS activity hardly moves, which validates CTP and dCTP’s strong inhibition on CTPS/dCTPS activity.

**Fig. 6.**
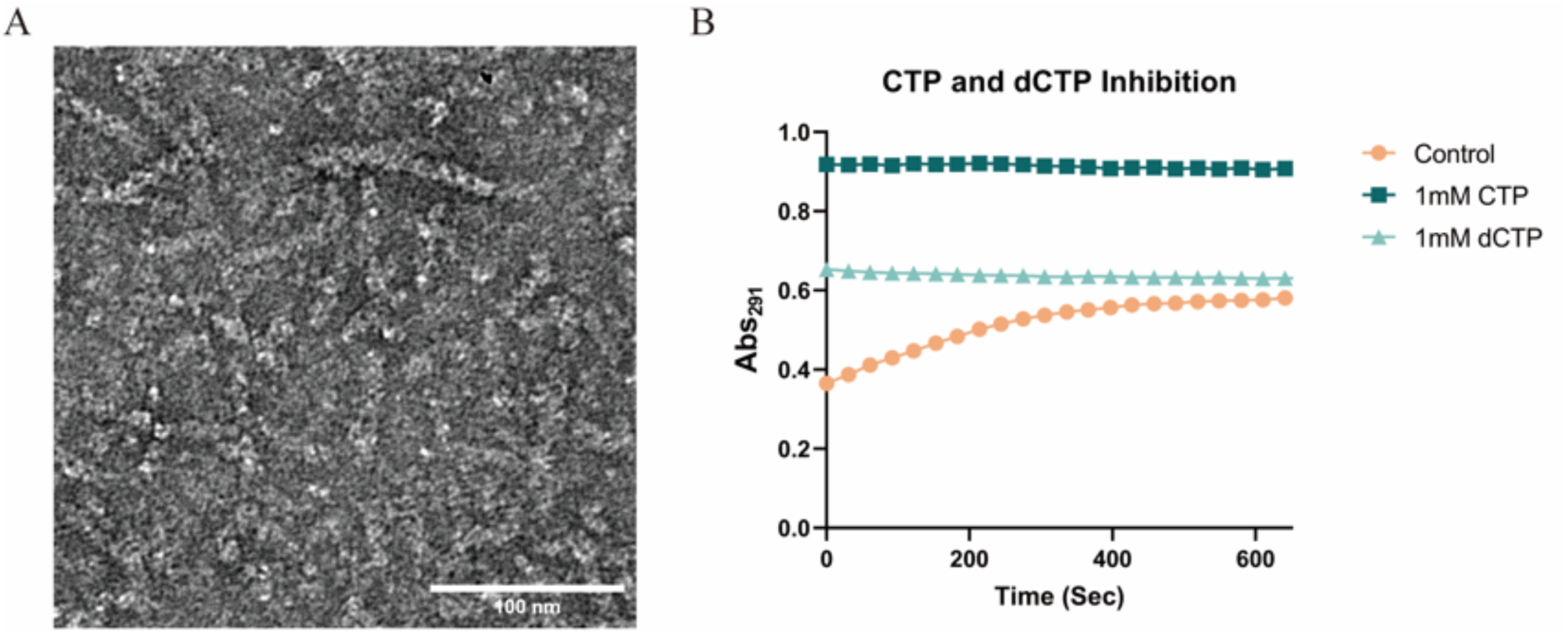
Effect of CTP and dCTP on CTPS/dCTPS. A, Negative-stain EM image of 3.2 μM CTPS mixed with 1 mM dCTP and 4 mM MgCl_2_. B, Absorbance readings of CTPS/dCTPS activity in control condition (0 mM CTP or dCTP), 1 mM CTP and 1 mM dCTP, respectively.

**Fig. 7.**
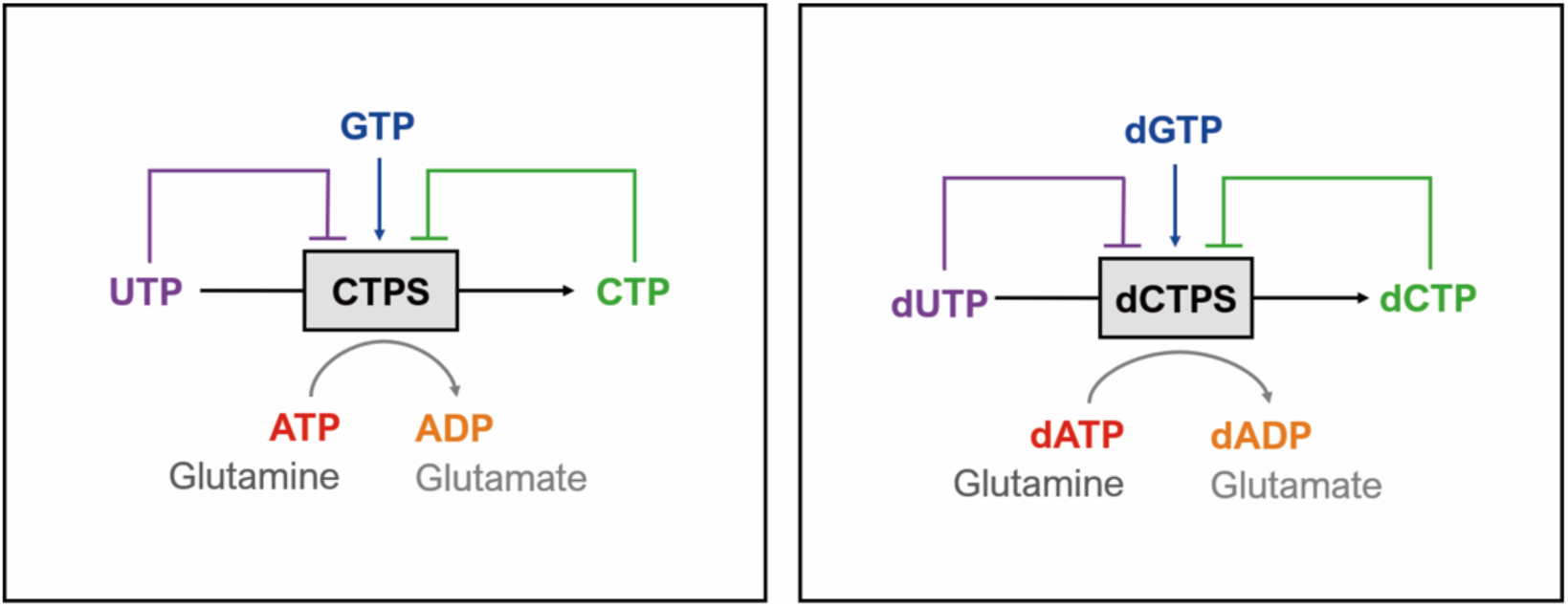
Catalytic diagram of bifunctional CTPS/dCTPS. It is well known that CTPS is the rate-limiting enzyme that catalyzes the last step in the de novo biosynthesis of CTP, and its catalytic process involves all four nucleotides that assemble RNA. The same enzyme catalyzes both CTP and dCTP production, highlighting its dual functions.

## Discussion

CTP/dCTP Synthase is the only enzyme that catalyzes the last step in de novo pyrimidine synthesis, yielding CTP, which is essential to all biological lives. CTPS/dCTPS is one of the most thoroughly and comprehensively studied complex metabolic enzymes, as it is generally considered as a classic example of biochemical regulation of key metabolic processes.

In 1999, Carman et al. reported that CTPS/dCTPS can tetramerize under deoxy-substrate conditions (using dATP, dUTP and dGTP as substrates). After more than 20 years, we finally revealed the structural basis for this novel catalytic mechanism that has been largely overlooked by the field in the past. Our work first reported that CTPS/dCTPS is capable of forming filaments using deoxyribonucleotides as substrates and regulators instead of using ribonucleotides, and we also provided structures of CTPS/dCTPS tetramers. However, in this paper, we did not investigate into how this molecular bifunctionality relate to cellular-scale filament formation termed cytoophidium. In the future, it would be interesting to study how filamentation on both molecular and cellular scale correspond with each other and how the relationship plays a role in CTP and dCTP catalysis.

In this paper, we advanced understanding of CTPS/dCTPS structure and orthogonally validated previous work through structural information. The relative conformational similarities between substrate and deoxy-substrate conditions may render dNTPs as a good substitute substrate for solving the dynamic structural states of CTPS/dCTPS and enzymes alike.

Our work provided structural basis for CTPS/dCTPS as a bifunctional enzyme that is capable of involving both ribonucleotides and deoxyribonucleotides in its catalysis. To fully appreciate the amazing properties of CTPS/dCTPS as a bifunctional enzyme, we should first note that standard CTPS/dCTPS catalysis involves all four ribonucleotides that assemble into RNA: ATP acts as its energy carrier; UTP acts as its substrate; GTP acts as its allosteric regulator; and CTP is its final product. With this in mind, when we observed that CTPS/dCTPS facilitates catalysis under deoxy-substrate conditions on subatomic level, we are fascinated by how flexible and compatible this enzyme is. why does CTPS/dCTPS accept both ribonucleotides and deoxyribonucleotides, two fundamental classes of biological macromolecules, in its catalytic process with such flexibility? Why keep both of these functions?

The first part of our explanation for the questions above is that bifunctionality allows CTPS/dCTPS to dynamically adjust its kinetic profile in order to adapt to the constantly changing environment. Our work showed that each deoxyribonucleotide’s kinetic parameters difference from those of ribonucleotides seem to correspond with the shift of conformations and ligand binding modes in deoxy-substrate conditions. In other words, kinetic similarity corresponds with structural similarity, and vice versa. This observation naturally follows that a range of binding mode possibilities exists for CTPS/dCTPS: if we only consider the three substrates, we now have eight distinctive combinations of either ribonucleotides or deoxyribonucleotides, each of which exhibits a different kinetic profile. Although we have not investigated into ligand selection preferences of CTPS/dCTPS when both ribonucleotides or deoxyribonucleotides are present, the inherent capacity for the enzyme to choose ligands and “customize” and fine tune its kinetic parameters is an amazing idea in itself.

Another major reason for CTPS/dCTPS bifunctionality is an alternative safeguard mechanism against Uracil Misincorporation. Uracil Misincorporation takes place when dUTP, instead of dTTP, is mistakenly incorporated into DNA. This phenomenon significantly damages DNA integrity, which leads to a variety of genetic disorders and is associated with high risk in cancer. According to current understanding of nucleotide metabolism, the only enzyme capable of degrading dUTP is dUTP diphosphatase (dUTPase), which converts dUTP into dUMP that is not cytotoxic. Our work now provides the structural basis for that CTPS/dCTPS is also capable of degrading dUTP into dCTP, which is not cytotoxic. This feature may have survived across time in order to prevent Uracil Misincorporation when deoxyuridine accumulates or when dUTPase fails to perform. Alternatively, CTPS/dCTPS may have always catalyzed dUTP along with dUTPase, but its coordinated behavior with dUTPase requires further research to be elucidated.

One might cast doubt on the claim that this flexible bifunctionality is an intelligent evolutionary design, but it would also be too dispirited to simply characterize this phenomenon as an anomalous feature arising from incomplete recognition of substrates during revolution. If we look at other nucleotide binding enzymes such as DNA Polymerase, they exhibit high-fidelity recognition of deoxyribonucleotides exactly for the purpose of not confusing between them with ribonucleotides. Moreover, as far as we currently know, there’s only one other enzyme that exhibits this bifunctionality: Nucleoside-diphosphate Kinase (NDPK). NDPK synthesizes more types of nucleotides, CTPS only synthesizes CTP/dCTP from UTP/dUTP. In light of such rarity, this bifunctionality is physiologically and metabolically significant.

As a rare exemplar of metabolic enzymes that lie at the crossroad between DNA and RNA catalysis, CTPS/dCTPS also shows good potential of broad applications in medicine, agriculture, and biological engineering. Through studying CTPS/dCTPS and its relationship to cellular filamentation, control of nucleotide pool balance and metabolic role, we may be able to apply this knowledge to tackle cancer and autoimmune diseases. Similarly, we may also engineer strains of plants that are more resistant to adverse conditions and engineer tools such as DNA/RNA sensors.

In the future, we would like to investigate whether CTPS/dCTPS cytoophidia, which are observable under confocal, are results of binding with NTP or dNTPs. Cytoophidia exist in all three domains of life and a variety of cell types. Furthermore, in fission yeast and homo sapiens cells, cytoophidia form not only in cytoplasm but also in nucleus. We wonder what the functional relationship between cytoophidia localization and its deoxy-substrate catalysis. Another question is: does cytoophidia formed under standard substrate conditions colocalize or communicate with cytoophidia formed under deoxy-substrate conditions?

In summary, our work advanced the field of inquiry into the bifunctionality of nucleotide metabolic enzymes such as CTPS/dCTPS. We extended previous work by providing structural basis for CTPS/dCTPS filamentation and catalysis under deoxy-substrate conditions. We believe that the bifunctionality of CTPS/dCTPS is connected with fundamental questions regarding the biophysics of filament self-assembly, flexible catalysis, and many other molecular and cellular mechanisms that are currently beyond our understanding. We also look forward to discovering more applications of CTPS/dCTPS and its bifunctionality in the future.

## Methods

### Protein expression and purification

A full-length *Drosophila melanogaster* CTPS sequence was constructed with a C-terminal 6XHis-tag and driven by T7 promoter. The plasmid was then transformed into Transetta (DE3) cells for expression. The cells were induced with 0.1 mM IPTG and incubated overnight at 16 °C. Cells were then pelleted by centrifugation at 4,000 rpm for 15 min followed by resuspension in cold lysis buffer (500 mM NaCl, 50 mM Tris-HCl (pH7.5), 20 mM imidazole, 1 mM PMSF, 7 µM leupeptin, 1.66 mM β-Me.) The cell lysate was then centrifuged (18,000 rpm) at 7 °C for 1 h. Supernatant was collected and fully mixed with equilibrated Ni-Agarose (Qiagen) for 1 h. Subsequently, Ni-Agarose was washed with a washing buffer (500 mM NaCl, 50 mM Tris-HCl (pH7.5), 64 mM imidazole, 5 mM β-Me.) The protein was then eluted with an elution buffer containing 500 mM NaCl, 50 mM Tris-HCl (pH7.5), and 266 mM imidazole. Hiload 16/600 Superdex 200 pg column and AKTA Pure (GE Healthcare) were used for further purification. Finally, dmCTPS was eluted with buffer containing 150 mM NaCl and 25 mM Tris-HCl (pH 8.0).

### Enzyme assays and analysis of kinetic data

UTP-dependent CTPS activity was determined by measuring the conversion of UTP to CTP through measuring increase in absorbance at 291 nm on a SpectraMax i3 (Molecular Devices) spectrophotometer. UTP and CTP have extinction coefficients of 182 and 1520 M^-1^ cm^-1^, respectively. Similarly, dUTP-dependent CTPS activity was determined by measuring the conversion of dUTP to dCTP through measuring increase in absorbance at 291nm; dUTP and dCTP have extinction coefficients of 137 and 1801 M^-1^ cm^-1^, respectively. The standard reaction mixture for UTP-dependent CTPS activity contained 12.5 mM Tris-HCl (pH 8.0), 75 mM NaCl, 10 mM MgCl_2_, 1 mM ATP, 1 mM UTP, 0.2 mM GTP, 10 mM Glutamine, and 2 μM CTPS. For dUTP-dependent CTPS activity, the reaction mixture composition is the same except for that 1 mM UTP is substituted with 1 mM dUTP. When conducting kinetic analysis, the appropriate component of the standard reaction mixture is changed to investigate how different concentrations of a certain component impacts the overall enzymatic activity. Enzyme concentration was determined by BCA. Kinetic data were analyzed using the GraphPad Prism 9.

### Negative stain microscopy

CTPS protein (3.175 μM) was dissolved in buffer containing 150 mM NaCl and 25 mM Tris-HCl (pH 8.0), 8 mM MgCl_2_, and 1 mM dCTP. After 30 min incubation at 37 degrees Celsius, 5 μL samples were loaded onto hydrophilic carbon-coated grids (400mech, Zhongjingkeyi Technology Co., China). Excess protein was blotted off from the grids before dying the grids. The grids were washed twice with 3.5 μL 0.5% uranium formate and then dyed a third time with 3.5 μL 0.5% uranium formate, which stayed on the grids for about 40 seconds before being blotted off. Imaging was performed on a 120 kV microscope (Talos L120C, ThermoFisher, USA) with an Eagle 4K X 4K CCD camera system (Ceta CMOS, ThermoFisher, USA). Images were acquired at 22000×, 36000x and 57000x magnification.

### Cryo-EM grid preparation and data collection

The Cryo-EM sample preparation condition is same to the negative staining. Amorphous alloy film (No. M024-Au300-R12/13) were hydrophilized for 40s before applying the 2.5µl sample solution. Then the grid was plunge-freezing by FEI Vitrobot (8 °C temperature, 3 s blotting time, −1 blot force). 3515 movies were collected by Gatan K3 summit camera with 22,500X magnification in super-resolution mode on a 300kv FEI Titan Krios electron microscope. Defocus is set to 0.8-1.6 μm, pixel size is 1.06Å. The total dose was 50 e−/Å2 subdivided into 40 frames at 2.8s exposure using SerialEM.

### Image processing

MOTIONCOR2 and CTFFIND4 were used to process the raw images by Relion 3.3 GUI. Particles were picking by the autopicker using reference generated by manual picking. After data cleaning by 2D and 3D classification, 108680 particles out of 1078812 particles were selected to the final reconstruction. CTF Refinement and Bayesian Polishing were implied to approve the resolution of map. Finally, a 3.53 Å map was obtained.

### 3D variability analysis and classification

Particles of the final reconstruction were exported to cryoSPARC V3.3 from RELION. Symmetry expansion was done by using D2 symmetry. And a protomer mask was used for the further processing. Three representative maps were selected from 3D variability analysis. Then they were used as templates for the 3D classification to generate three groups particles. Finally, maps of different states were generated by reconstruction only from each group of the particles.

### Model building and refinement

7WJ4 and 7DPT were used as initial model for Average state and Closed state model building. The manual fitting of models was done by using COOT. Then the coordinated is verified and tuned by Phenix.

## Acknowledgements

We thank Zixuan Wang and Xiaojie Bao for technical support. EM data were collected at the ShanghaiTech Cryo-EM Imaging Facility. We thank the Molecular and Cell Biology Core Facility (MCBCF) at the School of Life Science and Technology, ShanghaiTech University for providing technical support. This work was supported by grants from Ministry of Science and Technology of China (No. 2021YFA0804701-4), National Natural Science Foundation of China (No. 31771490), Shanghai Science and Technology Commission (No. 20JC1410500) and Medical Research Council for grants to J.L.L.

## Extended Data

**Supplementary Fig. S1.**
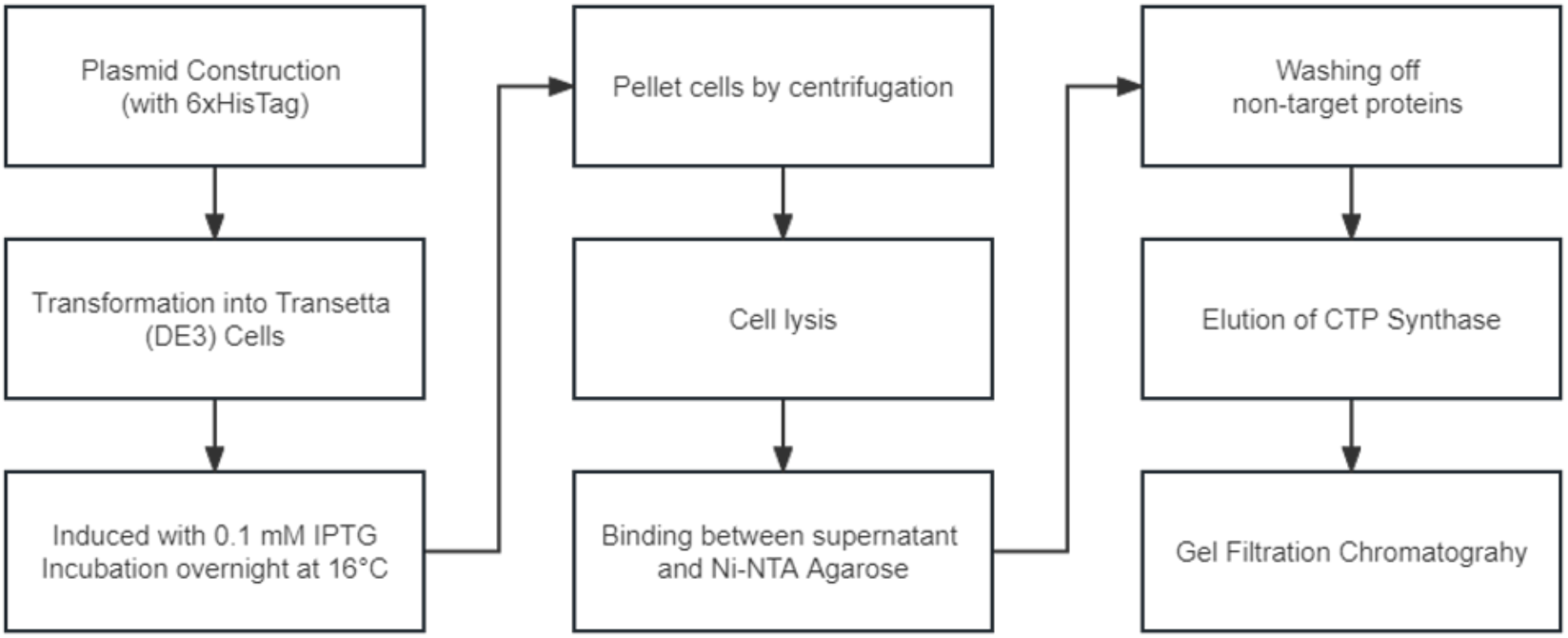
The flow chart of protein purification. Main procedure are indicated by box with annotation.

**Supplementary Fig. S2.**
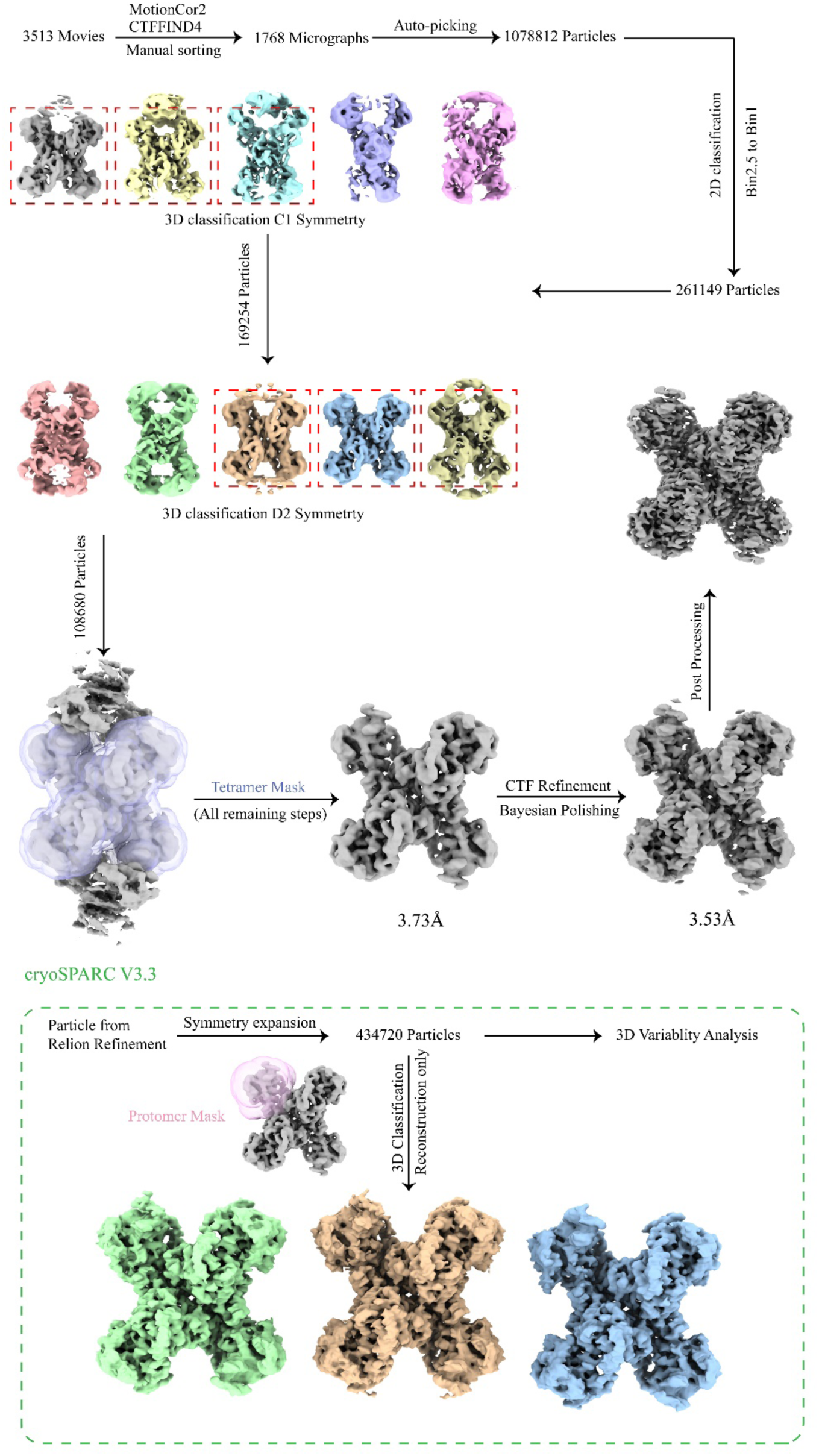
The work flow of data processing.

**Supplementary Fig. S3.**
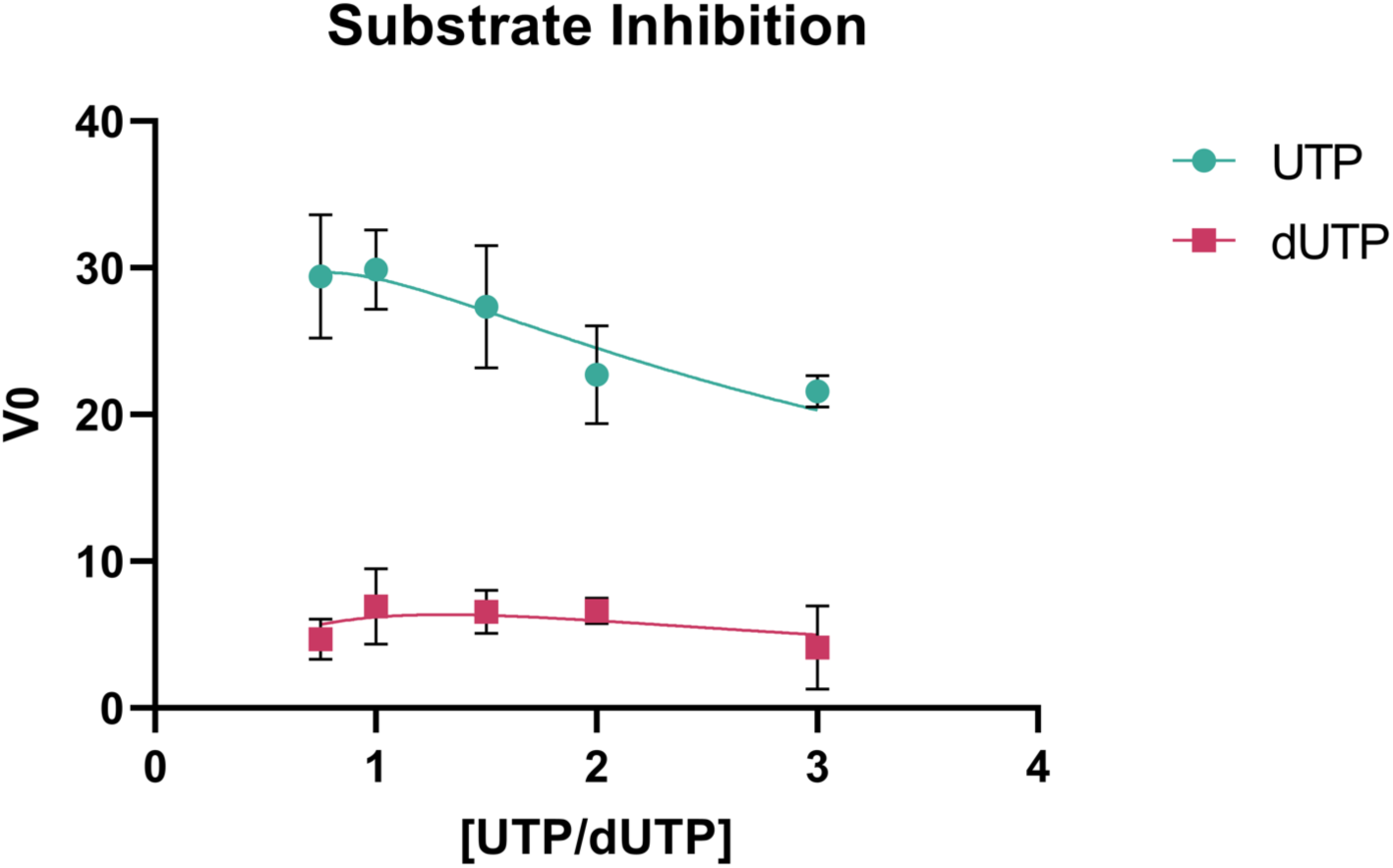
Substrate inhibition in CTPS/dCTPS activity using either UTP or dUTP as substrate.

**Supplementary Fig. S4.**
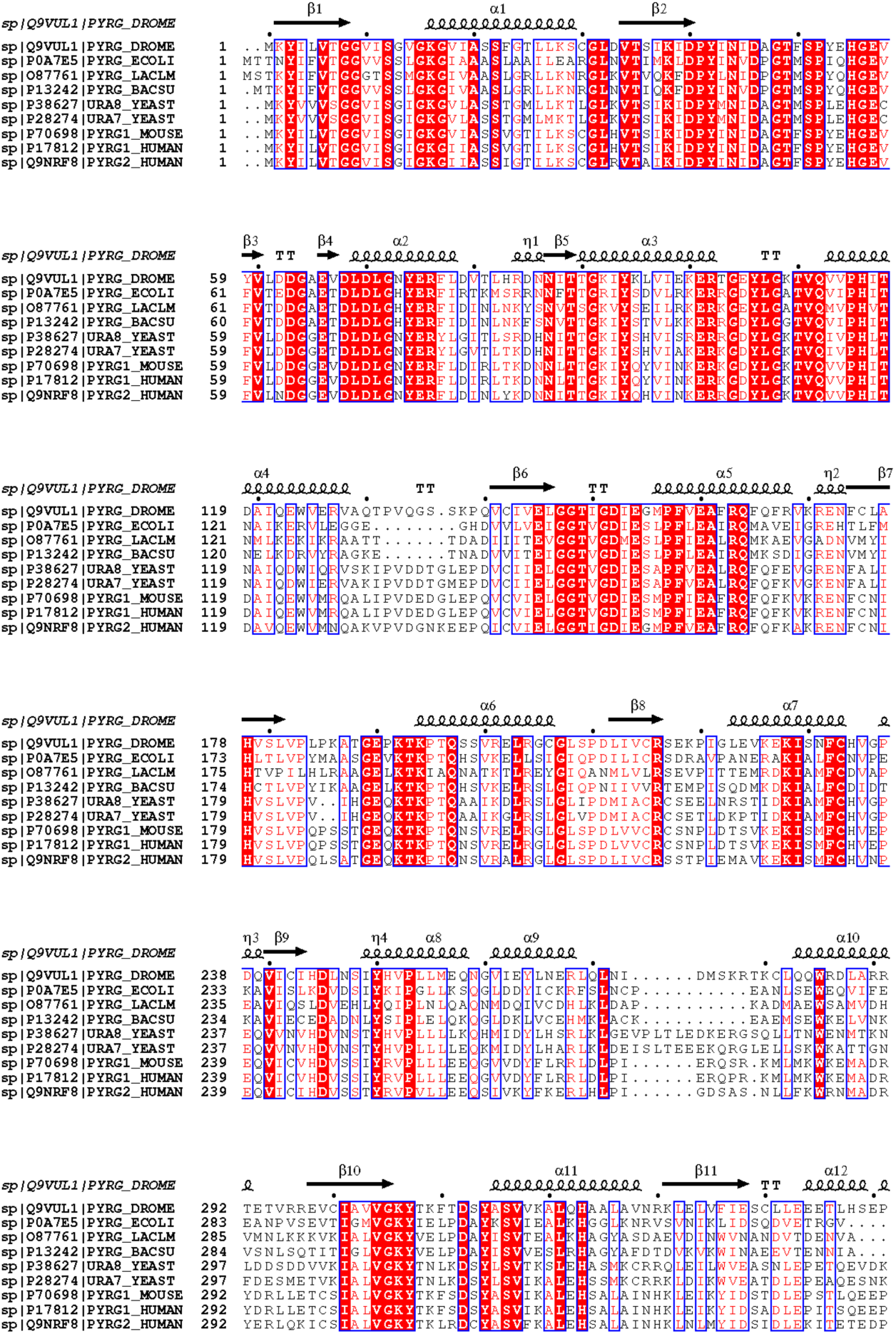
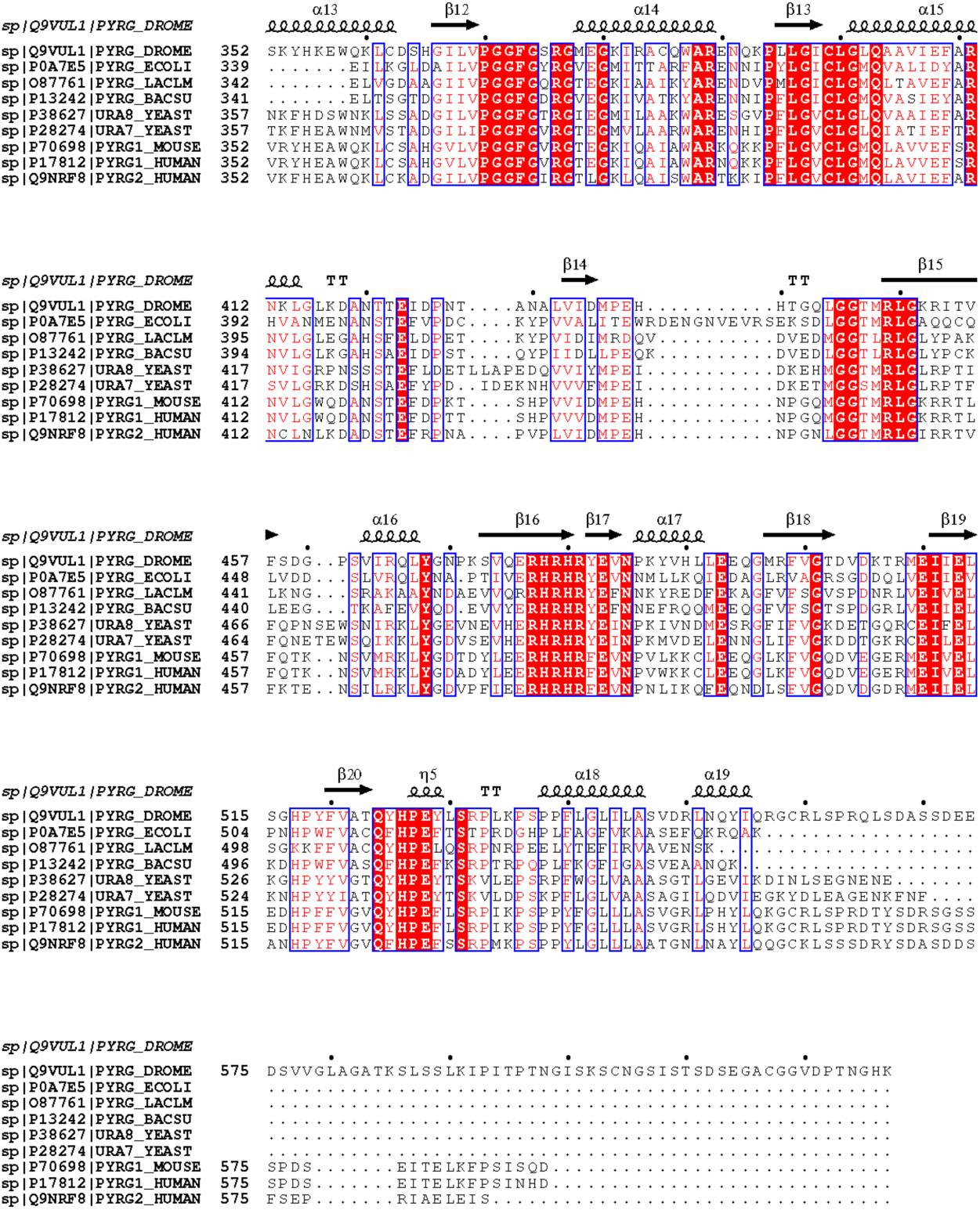
Sequence Alignment of CTPS/dCTPS among a variety of organisms.

**Table X.** Cryo-EM data collection and model refinement

**Supplementary Fig. S4. Sequence Alignment of CTPS/dCTPS among a variety of organisms.**
| Table X. Cryo-EM data collection and model refinement |  |  |  |  |
| --- | --- | --- | --- | --- |
| Model | Average model<br>(EMD-35103, PDB 8IOI) |  | Closed state model<br>(EMD-35102, PDB 8IOH) |  |
| Composition (#) |  |  |  |  |
| Chains | 12.00 |  | 7 |  |
| Atoms | 17772 (Hydrogens: 0) |  | 5102 (Hydrogens: 0) |  |
| Residues | Protein: 2224 Nucleotide: 0 |  | Protein: 635 Nucleotide: 0 |  |
| Water | 0.00 |  | 0 |  |
| Ligands | DUT: 4<br>DTP: 4<br>MG: 4 |  | DUT: 1<br>DTP: 1<br>MG: 2<br>DGT: 1 |  |
| Bonds (RMSD) |  |  |  |  |
| Length (?) (# > 4\|O) | 0.004 (0) |  | 0.005 (1) |  |
| Angles (\|a) (# > 4\|O) | 0.664 (12) |  | 0.845 (8) |  |
| MolProbity score | 2.15 |  | 2.27 |  |
| Clash score | 16.33 |  | 18.89 |  |
| Ramachandran plot (%) |  |  |  |  |
| Outliers | 0.18 |  | 0.16 |  |
| Allowed | 6.50 |  | 7.99 |  |
| Favored | 93.32 |  | 91.85 |  |
| Rama-Z (Ramachandran plot Z-score, RMSD) |  |  |  |  |
| whole (N = 2216) | -0.48 (0.18) |  | -0.84 (0.33) |  |
| helix (N = 888) | 0.70 (0.18) |  | 1.12 (0.30) |  |
| sheet (N = 372) | 0.02 (0.29) |  | -1.94 (0.78) |  |
| loop (N = 956) | -1.42 (0.20) |  | -2.24 (0.35) |  |
| Rotamer outliers (%) | 0.87 |  | 0 |  |
| C\|A outliers (%) | 0.00 |  | 0 |  |
| Peptide plane (%) |  |  |  |  |
| Cis proline/general | 3.6/0.0 |  | 6.1/0.0 |  |
| Twisted proline/general | 0.0/0.0 |  | 0.0/0.0 |  |
| CaBLAM outliers (%) | 2.54 |  | 2.17 |  |
| ADP (B-factors) |  |  |  |  |
| Iso/Aniso (#) | 17772/0 |  | 5102/0 |  |
| min/max/mean |  |  |  |  |
| Protein | 23.92/151.99/77.85 |  | 11.34/166.63/87.32 |  |
| Nucleotide | --- |  | --- |  |
| Ligand | 44.00/49.32/46.68 |  | 47.86/123.89/80.60 |  |
| Water | --- |  | --- |  |
| Occupancy |  |  |  |  |
| Mean | 1.00 |  | 1 |  |
| occ = 1 (%) | 100.00 |  | 100 |  |
| 0 < occ < 1 (%) | 0.00 |  | 0 |  |
| occ > 1 (%) | 0.00 |  | 0 |  |
| Data |  |  |  |  |
| Box |  |  |  |  |
| Lengths (?) | 138.73, 94.25, 123.90 |  | 90.10, 81.62, 89.04 |  |
| Angles (\|a) | 90.00, 90.00, 90.00 |  | 90.00, 90.00, 90.00 |  |
| Supplied Resolution (?) | 3.50 |  | 3.4 |  |
| Resolution Estimates (?) | Masked | Unmasked | Masked | Unmasked |
| d FSC (half maps; 0.143) | --- | --- | --- | --- |
| d 99 (full/half1/half2) | 3.7/---/--- | 3.7/---/--- | 3.5/---/--- | 3.4/---/--- |
| d model | 3.70 | 3.70 | 3.5 | 3.5 |
| d FSC model (0/0.143/0.5) | 3.1/3.2/3.7 | 3.2/3.2/3.7 | 3.2/3.5/4.2 | 3.3/3.5/7.1 |
| Map min/max/mean | -0.11/0.15/0.00 |  | -0.82/1.19/0.03 |  |
| Model vs. Data |  |  |  |  |
| CC (mask) | 0.74 |  | 0.72 |  |
| CC (box) | 0.72 |  | 0.48 |  |
| CC (peaks) | 0.65 |  | 0.23 |  |
| CC (volume) | 0.73 |  | 0.7 |  |
| Mean CC for ligands | 0.83 |  | 0.79 |  |

## References

1. Long, C.W. and A.B. Pardee, Cytidine Triphosphate Synthetase of *Escherichia coli* B: I. PURIFICATION AND KINETICS. Journal of Biological Chemistry, 1967. 242(20): p. 4715–4721.

2. Levitzki, A. and D.E. Koshland, Cytidine triphosphate synthetase. Covalent intermediates and mechanisms of action. Biochemistry, 1971. 10(18): p. 3365–3371.

3. von der Saal, W., P.M. Anderson, and J.J. Villafranca, Mechanistic investigations of Escherichia coli cytidine-5’-triphosphate synthetase. Detection of an intermediate by positional isotope exchange experiments. Journal of Biological Chemistry, 1985. 260(28): p. 14993–14997.

4. Ostrander, D.B., et al., Effect of CTP synthetase regulation by CTP on phospholipid synthesis in Saccharomyces cerevisiae. J Biol Chem, 1998. 273(30): p. 18992–9001.

5. Hatse, S., E. De Clercq, and J. Balzarini, Role of antimetabolites of purine and pyrimidine nucleotide metabolism in tumor cell differentiation. Biochem Pharmacol, 1999. 58(4): p. 539–55.

6. Williams, J.C., et al., Increased CTP synthetase activity in cancer cells. Nature, 1978. 271(5640): p. 71–3.

7. Kizaki, H., et al., Increased cytidine 5’-triphosphate synthetase activity in rat and human tumors. Cancer Res, 1980. 40(11): p. 3921–7.

8. Kang, G.J., et al., Cyclopentenylcytosine triphosphate. Formation and inhibition of CTP synthetase. J Biol Chem, 1989. 264(2): p. 713–8.

9. van den Berg, A.A., et al., Evidence for transformation-related increase in CTP synthetase activity in situ in human lymphoblastic leukemia. Eur J Biochem, 1993. 216(1): p. 161–7.

10. Agbaria, R., et al., Antiproliferative effects of cyclopentenyl cytosine (NSC 375575) in human glioblastoma cells. Oncol Res, 1997. 9(3): p. 111–8.

11. Lin, Y., et al., CTPS1 promotes malignant progression of triple-negative breast cancer with transcriptional activation by YBX1. J Transl Med, 2022. 20(1): p. 17.

12. Sun, Z., et al., Combined Inactivation of CTPS1 and ATR Is Synthetically Lethal to MYC-Overexpressing Cancer Cells. Cancer Res, 2022. 82(6): p. 1013–1024.

13. Martin, E., et al., CTP synthase 1 deficiency in humans reveals its central role in lymphocyte proliferation (vol 510, pg 288, 2014). Nature, 2014. 511(7509).

14. Lynch, E.M., et al., Structural basis for isoform-specific inhibition of human CTPS1. Proc Natl Acad Sci U S A, 2021. 118(40).

15. Fijolek, A., A. Hofer, and L. Thelander, Expression, purification, characterization, and in vivo targeting of trypanosome CTP synthetase for treatment of African sleeping sickness. J Biol Chem, 2007. 282(16): p. 11858–65.

16. De Clercq, E., Antiviral agents: characteristic activity spectrum depending on the molecular target with which they interact. Advances in virus research, 1993. 42: p. 1–55.

17. Mori, G., et al., Thiophenecarboxamide Derivatives Activated by EthA Kill Mycobacterium tuberculosis by Inhibiting the CTP Synthetase PyrG. Chem Biol, 2015. 22(7): p. 917–27.

18. Esposito, M., et al., A Phenotypic Based Target Screening Approach Delivers New Antitubercular CTP Synthetase Inhibitors. ACS Infect Dis, 2017. 3(6): p. 428–437.

19. Chiarelli, L.R., et al., A multitarget approach to drug discovery inhibiting Mycobacterium tuberculosis PyrG and PanK. Sci Rep, 2018. 8(1): p. 3187.

20. Levitzki, A. and D.E. Koshland, Jr., Role of an allosteric effector. Guanosine triphosphate activation in cytosine triphosphate synthetase. Biochemistry, 1972. 11(2): p. 241–6.

21. Robertson, J.G. and J.J. Villafranca, Characterization of metal ion activation and inhibition of CTP synthetase. Biochemistry, 1993. 32(14): p. 3769–77.

22. Bearne, S.L., O. Hekmat, and J.E. MacDonnell, Inhibition of Escherichia coli CTP synthase by glutamate gamma-semialdehyde and the role of the allosteric effector GTP in glutamine hydrolysis. Biochemical Journal, 2001. 356: p. 223–232.

23. Willemoës, M. and B.W. Sigurskjold, Steady-state kinetics of the glutaminase reaction of CTP synthase from Lactococcus lactis. The role of the allosteric activator GTP incoupling between glutamine hydrolysis and CTP synthesis. Eur J Biochem, 2002. 269(19): p. 4772–9.

24. Willemoes, M., et al., Lid L11 of the glutamine amidotransferase domain of CTP synthase mediates allosteric GTP activation of glutaminase activity. FEBS J, 2005. 272(3): p. 856–64.

25. Bakovic, M., M.D. Fullerton, and V. Michel, Metabolic and molecular aspects of ethanolamine phospholipid biosynthesis: the role of CTP:phosphoethanolamine cytidylyltransferase (Pcyt2). Biochem Cell Biol, 2007. 85(3): p. 283–300.

26. Lunn, F.A., J.E. MacDonnell, and S.L. Bearne, Structural requirements for the activation of Escherichia coli CTP synthase by the allosteric effector GTP are stringent, but requirements for inhibition are lax. J Biol Chem, 2008. 283(4): p. 2010–20.

27. Iyengar, A. and S.L. Bearne, An assay for cytidine 5’- triphosphate synthetase glutaminase activity using high performance liquid chromatography. Analytical Biochemistry, 2002. 308(2): p. 396–400.

28. Iyengar, A. and S.L. Bearne, Aspartate-107 and leucine-109 facilitate efficient coupling of glutamine hydrolysis to CTP synthesis by Escherichia coli CTP synthase. Biochem J, 2003. 369(Pt 3): p. 497–507.

29. Endrizzi, J.A., et al., Crystal structure of Escherichia coli cytidine triphosphate synthetase, a nucleotide-regulated glutamine amidotransferase/ATP-dependent amidoligase fusion protein and homologue of anticancer and antiparasitic drug targets. Biochemistry, 2004. 43(21): p. 6447–6463.

30. MacDonnell, J.E., F.A. Lunn, and S.L. Bearne, Inhibition of E. coli CTP synthase by the “positive” allosteric effector GTP. Biochimica Et Biophysica Acta-Proteins and Proteomics, 2004. 1699(1-2): p. 213–220.

31. Lunn, F.A. and S.L. Bearne, Alternative substrates for wild-type and L109A E. coli CTP synthases: kinetic evidence for a constricted ammonia tunnel. Eur J Biochem, 2004. 271(21): p. 4204–12.

32. McCluskey, G.D. and S.L. Bearne, “Pinching” the ammonia tunnel of CTP synthase unveils coordinated catalytic and allosteric-dependent control of ammonia passage. Biochimica Et Biophysica Acta-General Subjects, 2018. 1862(12): p. 2714–2727.

33. Zhou, X., et al., Structural basis for ligand binding modes of CTP synthase. Proceedings of the National Academy of Sciences of the United States of America, 2021. 118(30).

34. Weng, M. and H. Zalkin, Structural role for a conserved region in the CTP synthetase glutamine amide transfer domain. Journal of bacteriology, 1987. 169(7): p. 3023–3028.

35. Lewis, D.A. and J.J. Villafranca, Investigation of the mechanism of CTP synthetase using rapid quench and isotope partitioning methods. Biochemistry, 1989. 28(21): p. 8454–9.

36. Goto, M., et al., Crystal structures of CTP synthetase reveal ATP, UTP, and glutamine binding sites. Structure, 2004. 12(8): p. 1413–23.

37. Lauritsen, I., et al., Structure of the dimeric form of CTP synthase from Sulfolobus solfataricus. Acta Crystallogr Sect F Struct Biol Cryst Commun, 2011. 67(Pt 2): p. 201–8.

38. Zhou, X., et al., Drosophila CTP synthase can form distinct substrate-and product-bound filaments. Journal of Genetics and Genomics, 2019. 46(11): p. 537–545.

39. Lynch, E.M. and J.M. Kollman, Coupled structural transitions enable highly cooperative regulation of human CTPS2 filaments. Nat Struct Mol Biol, 2020. 27(1): p. 42–48.

40. Hansen, J.M., et al., Cryo-EM structures of CTP synthase filaments reveal mechanism of pH-sensitive assembly during budding yeast starvation. Elife, 2021. 10.

41. Park, T.-S., D.J. O’Brien, and G.M. Carman, Phosphorylation of CTP synthetase on Ser36, Ser330, Ser354, and Ser454 regulates the levels of CTP and phosphatidylcholine synthesis in Saccharomyces cerevisiae. Journal of Biological Chemistry, 2003. 278(23): p. 20785–20794.

42. Choi, M.-G. and G.M. Carman, Phosphorylation of human CTP synthetase 1 by protein kinase A: identification of Thr455 as a major site of phosphorylation. Journal of Biological Chemistry, 2007. 282(8): p. 5367–5377.

43. Choi, M.-G., T.-S. Park, and G.M. Carman, Phosphorylation of Saccharomyces cerevisiae CTP synthetase at Ser424 by protein kinases A and C regulates phosphatidylcholine synthesis by the CDP-choline pathway. Journal of Biological Chemistry, 2003. 278(26): p. 23610–23616.

44. Higgins, M.J., P.R. Graves, and L.M. Graves, Regulation of human cytidine triphosphate synthetase 1 by glycogen synthase kinase 3. Journal of Biological Chemistry, 2007. 282(40): p. 29493–29503.

45. Jia, F., C. Chi, and M. Han, Regulation of nucleotide metabolism and germline proliferation in response to nucleotide imbalance and genotoxic stresses by EndoU nuclease. Cell reports, 2020. 30(6): p. 1848–1861. e5.

46. Ingerson-Mahar, M., et al., The metabolic enzyme CTP synthase forms cytoskeletal filaments. Nat Cell Biol, 2010. 12(8): p. 739–46.

47. Liu, J.-L., Intracellular compartmentation of CTP synthase in Drosophila. Journal of Genetics and Genomics, 2010. 37(5): p. 281–296.

48. Noree, C., et al., Identification of novel filament-forming proteins in Saccharomyces cerevisiae and Drosophila melanogaster. Journal of Cell Biology, 2010. 190(4): p. 541–551.

49. Carcamo, W.C., et al., Induction of Cytoplasmic Rods and Rings Structures by Inhibition of the CTP and GTP Synthetic Pathway in Mammalian Cells. PLOS ONE, 2011. 6(12): p. e29690.

50. Chen, K., et al., Glutamine analogs promote cytoophidium assembly in human and Drosophila cells. J Genet Genomics, 2011. 38(9): p. 391–402.

51. Barry, R.M., et al., Large-scale filament formation inhibits the activity of CTP synthetase. Elife, 2014. 3: p. e03638.

52. Gou, K.M., et al., CTP synthase forms cytoophidia in the cytoplasm and nucleus. Exp Cell Res, 2014. 323(1): p. 242–253.

53. Petrovska, I., et al., Filament formation by metabolic enzymes is a specific adaptation to an advanced state of cellular starvation. Elife, 2014. 3.

54. Shen, Q.J., et al., Filamentation of Metabolic Enzymes in Saccharomyces cerevisiae. Journal of Genetics and Genomics, 2016. 43(6): p. 393–404.

55. Lynch, E.M., et al., Human CTP synthase filament structure reveals the active enzyme conformation. Nature structural & molecular biology, 2017. 24(6): p. 507–514.

56. Zhou, S., H. Xiang, and J.L. Liu, CTP synthase forms cytoophidia in archaea. J Genet Genomics, 2020. 47(4): p. 213–223.

57. Lynch, E.M., J.M. Kollman, and B.A. Webb, Filament formation by metabolic enzymes-A new twist on regulation. Current Opinion in Cell Biology, 2020. 66: p. 28–33.

58. Aye, Y., et al., Ribonucleotide reductase and cancer: biological mechanisms and targeted therapies. Oncogene, 2015. 34(16): p. 2011–21.

59. Pappas, A., T.S. Park, and G.M. Carman, Characterization of a novel dUTP-dependent activity of CTP synthetase from Saccharomyces cerevisiae. Biochemistry, 1999. 38(50): p. 16671–7.

